# Transient *grb10a* Knockdown Permanently Alters Growth, Cardiometabolic Phenotype and the Transcriptome in *Danio rerio*

**DOI:** 10.1101/2020.12.06.413633

**Authors:** Bridget L Evans, Terence Garner, Chiara De Leonibus, Oliver H Wearing, Holly A Shiels, Adam F L Hurlstone, Peter E Clayton, Adam Stevens

## Abstract

Embryonic growth trajectory is a risk factor for chronic metabolic and cardiovascular disorder. Grb10 is a negative regulator of the main pathways driving embryonic growth. This study investigates the long-term cardiometabolic consequences and transcriptomic profiles of transient disruption of grb10a expression in Danio rerio. Knockdown was associated with increased embryonic growth (+7%) and metabolic rate (+25%), and decreased heart rate (- 50%) in early life. Juvenile growth and respiratory rate were also elevated (+30% and 7-fold increase respectively). The transcriptome was permanently remodelled by this transient disruption, with dysregulation of multiple growth, cardiac, and metabolic pathways. Phenotypic alteration persisted into adulthood, resulting in a leaner body with elevated skeletal and cardiac muscle content and aerobic scope (43%). This study not only confirms for the first time that transient disruption of a single gene can result in permanent transcriptomic remodelling but correlates this remodelling with persistent alterations to the adult cardiometabolic phenotype.

## Introduction

The Foetal Origins of Adult Disease (FOAD) hypothesis proposes that diseases occurring in adulthood have their origins during development. While this hypothesis was first proposed in relation to coronary heart disease and foetal undernutrition in humans^1^, it is now accepted that a wide range of diseases have early developmental origins. In humans, small and large for gestational age (SGA, LGA) status are early risk factors for chronic metabolic and cardiovascular disorders, including type II diabetes (TIID), obesity, cardiac dysfunction, and hypertension^1–3^. Therefore, understanding the impact of altered embryonic growth trajectory on later life disease is important to identify at-risk individuals.

During embryonic development, temperature, circulating glucose levels, and oxygen availability serve as indicators of the mature, external environment^2–4^. Small changes to the phenotype occur in response to external cues to promote immediate survival in a process termed “developmental plasticity”. This is important for healthy development, though immediate survival can come at the expense of elevated disease risk in later life, particularly when there is mismatch between the developing and mature environments^3, 5^. While these correlations have been observed across a wide number of species, targeted, longitudinal in-vivo studies to elucidate the developmental origins of health and disease, and the pathways involved, are lacking.

Zebrafish are an ideal model organism for both longitudinal and developmental study owing to a rapid generation time, large clutch sizes, and ease of access to embryos. As in mammals, embryonic growth in zebrafish is primarily driven by the insulin/insulin like growth factor (Ins/IGF) signaling pathway^6^. Circulating insulin and IGF levels give an indication of the caloric and nutrient condition of the mature environment, allowing modulation of developmental rate to match the prevailing conditions and improve survival prospects.

Growth factor receptor bound protein 10 (GRB10) is a negative regulator of the Insulin/IGF signaling pathway. GRB10 downregulates the growth response, promoting a switch from glucose to fat metabolism and halting cell cycle progression^7–9^. GRB10 expression limits placental growth and efficiency^10^ and correlates with small body size ^11^ in mammals. In humans, GWAS show *GRB10* is associated with TIID^12^, and *GRB10* copy number variation is associated with Silver Russell Syndrome^13^, a rare growth disorder typically characterised by intrauterine growth restriction, SGA status, hypoglycemia, poor muscle development, and increased fat deposition. The role of GRB10 in the regulation of human growth is notably associated with response to recombinant human growth hormone in children with growth hormone deficiency, where lower GRB10 expression correlates with a greater response^14, 15^. Variability in Grb10 expression is also linked to the dramatic range of body sizes observed between cetaceans^16^. Average daily mass gain is elevated in beef cattle with a *grb10* associated deletion^17^, and *grb10* SNPs impact muscle and lipid mass, angularity, and body conditioning score^18^. Global *grb10* knockout in mice correlates with a “leaner” phenotype, including elevated muscle and reduced lipid mass, and insulin sensitivity^19, 20^. Therefore, understanding the impact transient disruption to early-life growth trajectory has on mature organism size, average daily gain, and lipid to muscle ratio may provide the groundwork for boosting meat yield and quality, necessary to match the growing global demand for protein. In this study, *grb10a* expression was transiently supressed in wild-type zebrafish (*Danio rerio*) embryos by antisense oligonucleotide directed blocking of mRNA splicing, resulting in increased insulin/IGF signalling during development. The importance of *grb10a* as a coordinator of growth, metabolism, and cardiac health was assessed over the first 5 days post fertilisation (dpf). The impact of early-life growth disruption on the transcriptomic landscape was also investigated, together with the lasting impact on later-life body morphology, metabolism, and cardiac phenotype, linking together the distinct pathways commonly associated with the FOAD hypothesis.

## Materials and Methods

### Zebrafish Husbandry

AB zebrafish were maintained under standard conditions (≈28 °C; 14/10 h light/dark cycle; < 5 fish per litre) within the Biological Services Unit of The University of Manchester. Regulated procedures received ethical approval and were performed under a Home Office Licence (PPL P005EFE9F9). To generate embryos, breeding pairs of similar ages were selected at a ratio of 1 male to 1 female and fasted overnight in breeding tanks. Dividers were removed at the start of the following light cycle and embryos were collected after 20 minutes of free breeding. Embryos were kept at a stocking density of < 50 per petri dish and raised in embryo water (Instant Ocean salt 60 µg/mL) up to 5 dpf and transferred to the main aquarium.

### Transient Knockdown of *grb10a* Expression

Morpholino-modified antisense oligonucleotide knockdown (KD) of grb10a was validated in accordance with current guidelines for morpholino use in zebrafish^21^. Morpholinos targeting exon three (e3i3) and four (e4i4) were designed by and obtained from Gene Tools, LLC (Philomath, OR, USA) along with a standard control (SC) oligonucleotide targeting human β- globin, used to control for microinjection (sequences - *Table 1,* microinjection solutions *- Table 2*^21^). Phenol red and nCerulean (nuclear-targeting blue fluorescent protein) mRNA were included to ensure successful injection. Embryos received a single injection into the yolk directly below the cell mass at the single-cell stage, as per established methods^21^. Embryos were screened for fluorescence at 48 hours post-fertilisation (hpf) to ensure constitutive and even uptake of the injection material. Non-uniformly or weakly-fluorescent embryos were removed.

**Table 1.**
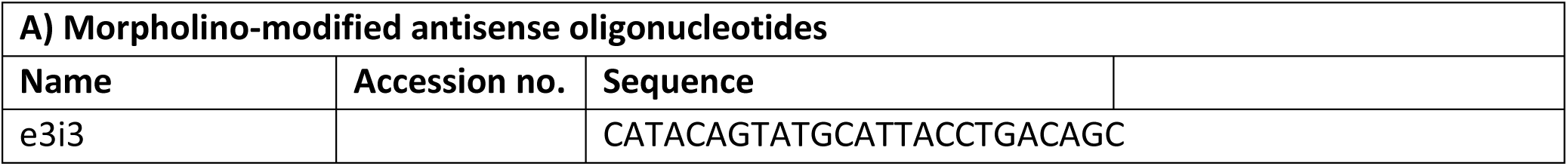
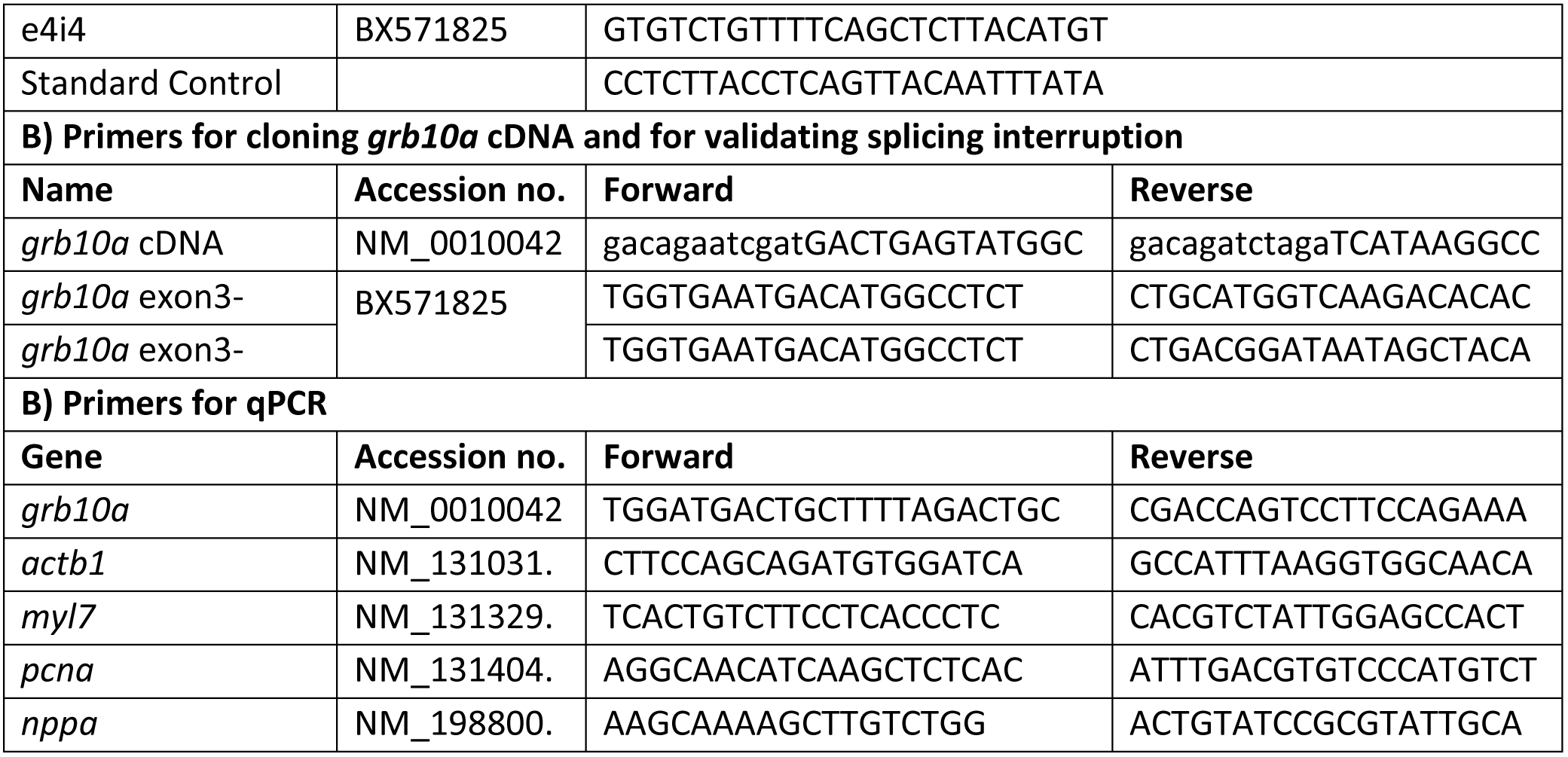
Oligonucleotide sequences.

**Table 2.** Microinjection Solutions.

| Injection | Oligonucleotide | nCerulean | Phenol Red |
| --- | --- | --- | --- |
| Standard Control (5 | 0.5 mM human beta globin | 50 ng/μl | 1 μl |
| *grb10a* KD (5 μl) | 0.5 mM e3i3 or e4i4 morpholino | 50 ng/μl | 1 μl |
| *grb10a* OE (5 μl) | 500 ng/μl *grb10a* RNA | 50 ng/μl | 1 μl |

### Validation of *grb10a* Knockdown

Primer sequences, outlined in Table 1, were designed using SnapGene® (GSL Biotech, San Diego, CA, USA) and synthesised by Thermo Fisher Scientific (Waltham, MA, USA). Specificity was confirmed using Primer BLAST^22^. To confirm antisense oligonucleotide activity, RNA was extracted from pooled zebrafish embryos at 24, 48, 72, 96, and 120 hpf (n=3, 5 embryos per pool). Extraction was performed using QIAGEN RNeasy lipid extraction kit according to the manufacturer’s instructions, and cDNA was generated by reverse transcription using the ProtoScript® II First Strand cDNA Synthesis Kit (NEB). cDNA was amplified with primers flanking each splice site by Taq polymerase (NEB, Hitchin, UK) (thermocycling parameters outlined in Table 3). β-actin (*actb1*) was used as a positive control for cDNA integrity.

**Table 3.**
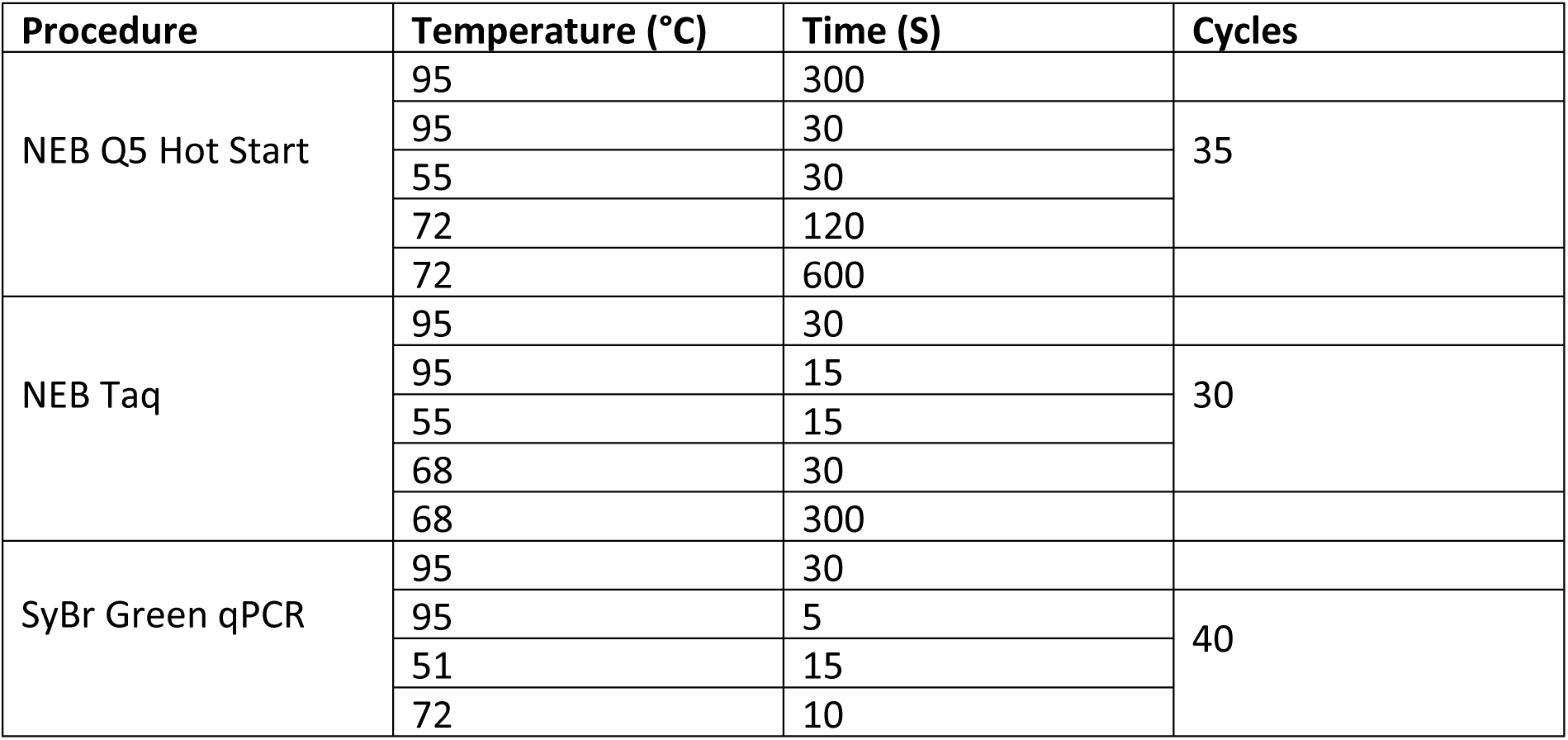
Thermocycling Parameters.

### Analysing Downstream Signalling by Western Blot

96 hpf embryos were deyolked in Ringer’s Buffer to limit background interference (n=15, performed in triplicate). Embryos were resuspended in 100 μl RIPA buffer (150 mM NaCl, 1% Nonident P-40, 0.5% Sodium deoxycholate, 0.1% SDS, 25 mM Tris pH 7.4) containing protease and phosphatase inhibitors, and homogenised. Samples were incubated on ice for 30 minutes, clarified by centrifugation at 4 °C, and denatured at 98 °C for 5 minutes in Laemmli buffer (2% SDS, 10% glycerol, 60 mM Tris-Cl, 0.01% bromophenol blue, 0.1% β-Mercaptoethanol). Proteins were separated by 10% SDS acrylamide gel electrophoresis and transferred using established methods. The transfer membrane was incubated in blocking buffer (3% BSA in TBS-T) for one hour, primary antibody (Table 4) at 4 °C under constant agitation overnight, and secondary antibody (Table 4) for one hour at room temperature. The membrane was covered with ECL Western Blotting Substrate (Promega, Southampton, UK) and imaged immediately. Protein expression was quantified from band intensity in ImageJ^23^.

**Table 4.** Western Blot Antibodies.

| Target | Species | Supplier | Dilution |
| --- | --- | --- | --- |
| AKT | Mouse | Cell Signalling #2920 | 1:1000 |
| p-AKT | Rabbit | Cell Signalling - #5831 | 1:1000 |
| S6 | Mouse | Cell Signalling - #2317 | 1:1000 |
| p-S6 | Rabbit | Cell Signalling - #2215 | 1:1000 |
| Anti-Mouse HRP linked<br>IgG | Horse | Cell Signalling - #7076 | 1:5000 |
| Anti-Rabbit HRP linked<br>IgG | Goat | Cell Signalling - #7074 | 1:5000 |

### *grb10a* mRNA for Overexpression and Rescue

Total RNA was extracted, and cDNA generated from a pool of 96 hpf embryos, as previously described. *Grb10a* was amplified from cDNA by high specificity PCR (NEB Q5 Hot Start) with primers flanking open reading frame (Table 1). The product was purified using QIAGEN Quick Gel Extraction, blunt ligated into pCR-Blunt II-TOPO (Thermo Fisher Scientific) and transformed into *E. coli* following an established protocol. The purified plasmid was Sanger sequenced to confirm successful cloning. The insert was liberated by digestion with *Cla I* and *Xba I* restriction enzymes (NEB) and subcloned into pCS2+^24^. Capped RNA was generated using the mMESSAGE mMACHINE® SP6 Transcription Kit (Thermo Fisher Scientific) according to the manufacturer’s instructions. RNA was purified by MEGAclear™ Transcription Clean- Up. 1 μl of the purified RNA was analysed by gel electrophoresis to confirm amplification and structural integrity.

### Embryonic Physiological and Metabolic Measurements

Whole body length, the longest straight-line distance between the snout and tip of the notochord, and yolk area measurements were taken at 24-hour intervals from 48 to 120 hpf. Embryos were dechorionated and acclimatised for ≥ one hour before imaging. Images were imported into ImageJ, which was calibrated with an image of a graticule of known size. Data were imported into GraphPad Prism version 7.00 for Windows (GraphPad Software, La Jolla, CA, USA, www.graphpad.com).

To investigate embryonic cardiac phenotype, embryos were sedated using an anaesthetic concentration known to have no impact on cardiac function (0.04% MS-222 solution^25^). Heart beats were counted over a 20 second period and converted to beats per minute (bpm).

A Glucose Uptake-Glo TM Assay (Promega) was performed on 96 hpf zebrafish to detect differences in metabolic rate. Individual embryos (n = 5 per treatment) were injected with 1 mM 2-deoxyglucose-6-phosphate directly into the yolk and allowed to recover for 30 minutes. Embryos were processed according to an established protocol^26^. Luminescence was measured by plate reader with 8 readings per well. Readings were adjusted for background luminescence.

### Quantitative PCR

QPCR primers amplifying *grb10a* and markers of cardiac dysfunction were designed (Table 1) and their efficiency validated^27^. RNA was extracted from pooled (n=10) embryonic zebrafish and pooled (n=3) adult (> 1 year) heart samples by QIAGEN RNEasy Lipid Tissue Extraction kit according to the manufacturer’s instructions. Samples were repeated in triplicate and tested for gene expression by qPCR (Applied Biosystems Power SyBr Green) (Table 3). Relative fold change in gene expression was calculated using the ΔΔCt method according to the following equation:

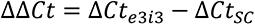

Where

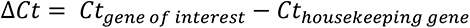

and

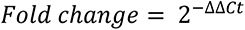

Data were imported into GraphPad and unpaired t-tests were performed on ΔCt values to assess the statistical difference between samples.

### Transcriptomic Analysis

Age-associated changes in gene expression were assessed by computational analysis of transcriptomic data. Pooled samples (n=5) were generated from each treatment group (SC, KD) at 5, 10, 15, 20, and 30 dpf, with three repeats per sample. Zebrafish were culled under terminal anaesthesia, and RNA was extracted from tissue anterior to the gills.

Transcriptomic data were generated using Affymetrix Zebgene 1.0st arrays. Data for all 75212 gene probes were imported into Qlucore Omics Explorer 2.2 (Lund, Sweden) as .cel files and normalised using the robust multi-array average (RMA) approach with a gene level summary. Zebrafish gene identities were assigned using Affymetrix gene definitions. Human orthologues (GRCh 38) were mapped using the *biomaRt* R-package^28^. A workflow pipeline of the transcriptomic analyses is outlined in *Supplementary Figure 1*.

Unsupervised analysis of gene expression by age group was conducted by generating hierarchically clustered heat maps. Standard deviation filtering (standard deviation of specific gene expression divided by maximum gene standard deviation [*S*/*S*_max_]) was performed on the dataset to remove genes with low variance, as these were unlikely to be informative. Projection scores^29^ were used to determine the threshold for filtering by calculating the maximum separation in principal component analysis (PCA).

### Hypernetwork Modelling

Hypernetwork analysis was performed to investigate higher order interactions between target genes, a general model of which is outlined in *Supplementary Figure 1*^30^. All analyses were performed in R (version 3.4.2). Pearson’s correlation coefficients (*r*) were calculated between age-associated genes identified by unsupervised analysis (*g,* KD = 119, SC = 297) and the rest of the transcriptome (*g^c^*, KD = 75093, SC = 74915). R-values were binarized to generate the incidence matrix of the hypernetwork (*M*). Positive and negative correlations greater than ±1 standard deviation (sd) from the mean of the R-values were assigned as ‘1’ (i.e., present) and values closer to zero were assigned ‘0’. In this way, each element (∈) of *g* can be described as:

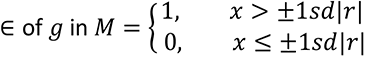

The resulting binary incidence matrix (*M*) was multiplied by its transpose (*M^t^*) to generate the adjacency matrix of the hypernetwork (*M*.*M^t^*) which quantifies the shared correlations between any pair of genes (g). This hypernetwork adjacency matrix represents the higher order interactions between pairs of genes in a manner not captured by traditional transcriptomic approaches. As a measurement of co-ordination, hypernetworks have been suggested to model functional relationships^30^. Coordination between age-related genes and the rest of transcriptome was investigated by interrogating the incidence matrix of the hypernetwork and identifying a subset of edges and nodes which could form a complete subgraph. This represents the subset of the transcriptome (⊂ *g^c^*) showing correlated expression with all the transcripts from the hypernetwork central cluster (⊂ *g*).

### Quantification of Network Topology

All subsequent analyses focused on genes defined as the central cluster of the hypernetwork (86 genes in SC fish, 67 in KD) of 20-30 dpf zebrafish. Correlation networks model functional relationships within gene networks^32^ and allow identification of clusters of highly connected genes^33^. Quantification of hypernetwork properties (connectivity and entropy) was performed on the hypernetwork (*M*.*M*^t^), where connectivity is the sum of connections shared by each element of the network (*g*) and entropy is the degree of disorder within the distribution of shared correlations. Entropy is positively correlated with the cellular differentiation potential^34^, where a high entropy indicates an earlier cell lineage and multiple potential signalling pathways, and low entropy indicates a more specific function. Entropy has also been used as an index of regularity and patterning^35^. Entropy was measured using R package BioQC^36^.

To identify the impact of age-association on the transcriptome and compare the effect between the SC and KD data sets, connectivity and entropy were calculated and averaged for 1000 hypernetworks of genes randomly selected from each dataset (having no age- association). The difference in connectivity and entropy that age-association conveyed on the transcriptome could then be determined by comparing the age-associated datasets with their corresponding random datasets. As entropy scales proportionally with the size of a set, entropy was normalised for each dataset, generating a proportional measure of the maximum entropy possible for a given set size (ranging from 0 - 1), calculated by dividing the measured entropy by log(n) where n is the set size.

### Gene Ontology

Gene set enrichment analysis (GSEA) was performed^37^ to associate gene expression with biological processes. Genes were mapped to human orthologues using Qlucore and GSEA was performed to rank genes by age group associated ANOVA p-values. Additional GSEA was carried out through Webgestalt^38^ using genes ranked by R-value, derived from a rank regression analysis of gene expression against age. Over-representation analysis (ORA) was used to identify gene ontology associated with unranked gene sets (Webgestalt). All gene ontology analysis used the GO Biological Process Ontology gene list^37, 39^.

### Hypernetwork Modelling of Ontology

Pathways identified by GSEA were modelled using a hypernetwork approach to investigate the association between each pathway and the two treatment groups. Hypernetworks were generated separately for SC and KD fish using human genes associated with each pathway^40^. Pathways with fewer than 15 associated genes were removed, and all remaining gene sets were converted to zebrafish homologues using the Ensembl database (release 104)^41^, queried using BiomaRt for R^28, 42^.

Hypernetworks were generated on 10 genes from each pathway, iterated 1000 times. Hypernetwork entropy was assessed on each iteration. A Bayesian approach was used to model the entropy distributions for each pathway to identify differences between SC and KD. This was performed using Bayesian generalized linear modelling via the r package rstanarm^43, 44^. Differences between SC and KD entropy distributions were calculated as a β value and significance was assigned to pathways for which the 89% credible interval of the beta values did not include 0, as per established methods^45^.

### Stop-Flow Respirometry

Individual 30 dpf zebrafish were placed into one of four stop-flow respirometry chambers (volume 2 ml) and allowed to acclimate at 28 °C for > one hour. Optical oxygen sensors paired with oxygen sensor spots (Pyroscience, Aachen, Germany) were used to measure oxygen saturation within the chambers and recorded using a FireStingO2 Fiber-optic oxygen and temperature meter, simultaneously recording and maintaining temperature at 28 ± 0.3 °C. Probes were calibrated according to the manufacturer’s instructions. Chambers were randomised per trial, with one chamber left empty to correct for background bacterial respiration. Chambers were refreshed immediately prior to the experiment until oxygen saturation measured 100%. Oxygen consumption curves were recorded in triplicate with five trials per individual. Chambers were manually refreshed when oxygen saturation reached 80%. To calculate the rate of oxygen consumption, linear regression in Microsoft Excel was used to calculate the change in oxygen saturation during each trial. This was normalised against the dry mass of the subject, length of the trial, and volume of the respirometry chamber according to the following equation:

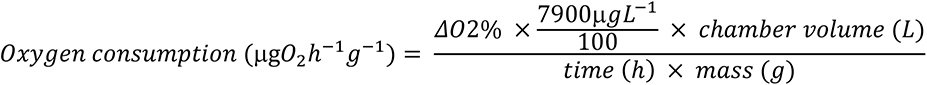

Where 7900 μgL^-1^ is equivalent to 100% oxygen saturation at 28 °C.

A closed-circuit stop-flow respirometer was used to investigate aerobic scope in adult zebrafish (18 months). Sealed chambers (70 ml) were combined with recirculation loops containing an oxygen flow-through cell (Pyroscience, Aachen, Germany), paired with an optical oxygen sensor (Pyroscience, Aachen, Germany), calibrated according to the manufacturer’s instructions. A stop-flow pump, controlled by a Cleware USB-Switch (Cleware GmbH, Germany) programmable switch and AquaResp v.3 software (AquaResp, v3, Python 3.6^46^) was incorporated into the circuit to automatically refresh the water in the chambers after each trial (60 s flush, 30 s wait, 300 s measure). To maintain a constant temperature of 28 °C ± 0.3 °C, the respirometry system was immersed in a recirculation chamber under constant aeration. Oxygen saturation and water temperature were recorded as previously described. Regression curves were automatically generated by AquaResp, and oxygen consumption was extracted and normalised against the length of the trial and volume of the respirometry chamber. Zebrafish were manually stressed for 2 minutes immediately prior to the start of the first trial. Maximum oxygen consumption was identified as the trial with the greatest difference in oxygen saturation. Standard metabolic rate was calculated as the mean of the lowest 10% of the trials^47^. Aerobic scope was calculated as the difference between the maximum oxygen consumption and standard metabolic rate. Data are presented as individual data points alongside the mean and SEM.

### Glucose Tolerance and Insulin Sensitivity

Adult zebrafish (18 months) were fasted overnight and allocated to either glucose tolerance testing or insulin sensitivity testing. All blood samples were acquired following the protocol outlined by Zhang et al.^48^. Mass, body length, and fasting blood glucose were measured immediately prior to the start of the protocol.

Fish were anaesthetised in 0.02% MS-222, placed on their side and patted dry. A single IP injection of glucose (0.5 mg glucose/g) or glucose and insulin (0.5 mg glucose/g and 0.0075 U insulin/g) was performed before recovery in 28 °C system water^49, 50^. Blood samples were taken at 30- and 120-minutes post-injection (glucose tolerance) or 30- and 60-minutes post injection (insulin sensitivity). Blood samples were immediately tested for blood glucose concentration (Sinocare Safe AQ blood glucose monitor). Individuals were culled in MS-222 during the final blood draw. Data are presented as the mean and the SEM (glucose tolerance n = 10, insulin sensitivity n = 8-12).

### Adult Physiological Measurements

To assess the end-stage body morphology induced by *grb10a* KD, dry mass and body length were measured in adult (18 month) zebrafish and Fulton’s condition factor was calculated. Body length was measured as the greatest straight-line distance between the snout and the end of the tail. The caudal fin was not included as fin length can be influenced by factors such as damage or variation between strains. Fulton’s condition factor was calculated according to the formula^51^:

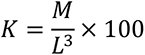

Where M = mass in grams and L = length in centimetres. Higher K values indicate thicker, more rounded bodies. Data are presented as the mean ± SEM (n = 21-34).

### Histology

Whole adult zebrafish (18 months) were embedded longitudinally in paraffin wax. 5 µm sagittal sections were taken from each tissue at a consistent depth and stained with Masson’s Trichrome (IHC World Masson’s Trichrome Staining Protocol for Collagen Fibres, Woodstock, MD, USA) to differentiate skeletal muscle (red), connective tissue (blue), and nuclei (black). Slides were scanned and visualised at 20x magnification (3D Histech CaseViewer v2.4.0.119028, Budapest, Hungary).

Skeletal muscle measurements were performed on a site lateral to the dorsal fin. The perpendicular width of individual muscle fibres was recorded and are presented as the mean of five measurements of each muscle fibre (ten fibres from five individuals, 50 fibres total).

Red blood cells were digitally removed from images of the ventricle before importing into ImageJ. The ratio of compacta to spongiosa was calculated by measuring the area of each tissue type. Data are presented as the ratio of compacta to spongiosa as a percentage. Cardiac tissue density was calculated by restricting the region of interest to the boundary of the ventricle and calculating the total number of pixels in the image. Threshold_Colour was used to threshold the images, which were converted to 8-bit black and white images. Voxel_Counter.class was used to calculate the number of black pixels in the image.

### Statistical Tests

For transcriptomic analyses, rank regression (least squares method) was used to generate the most appropriate linear model for each probe (the smallest degree of variance over the sample). Multi group analysis of variance (ANOVA) was used to associate each gene ID with time dependent gene expression. Wilcoxon rank sum test (ggpubR package for R^52^) was used to test for differences in network topology. False discovery rate (FDR) adjustment was made using the Benjamini-Hochberg method and applied to the gene ontology analysis^53^.

All data were ROUT tested^54^ for outliers and subject to D’Agostino and Pearson normality tests. All comparisons between SC and KDs were performed using unpaired t-tests. Comparisons of multiple groups were performed using one-way ANOVAs. Post-hoc power calculations were performed to confirm sample sizes were sufficient, where α = 0.05.

## Results

### Knockdown of *grb10a* Expression by Splice-Blocking Antisense Oligonucleotide

As *grb10* has been linked to embryonic growth trajectory, expression was examined over the first 120 hpf. QPCR analysis of *grb10a* expression at 24-hour intervals revealed a strong upregulation at 48 hpf (*Figure 1a*). Expression of the *grb10* paralogue, *grb10b*, was not detectable at any time point.

**Figure 1:**
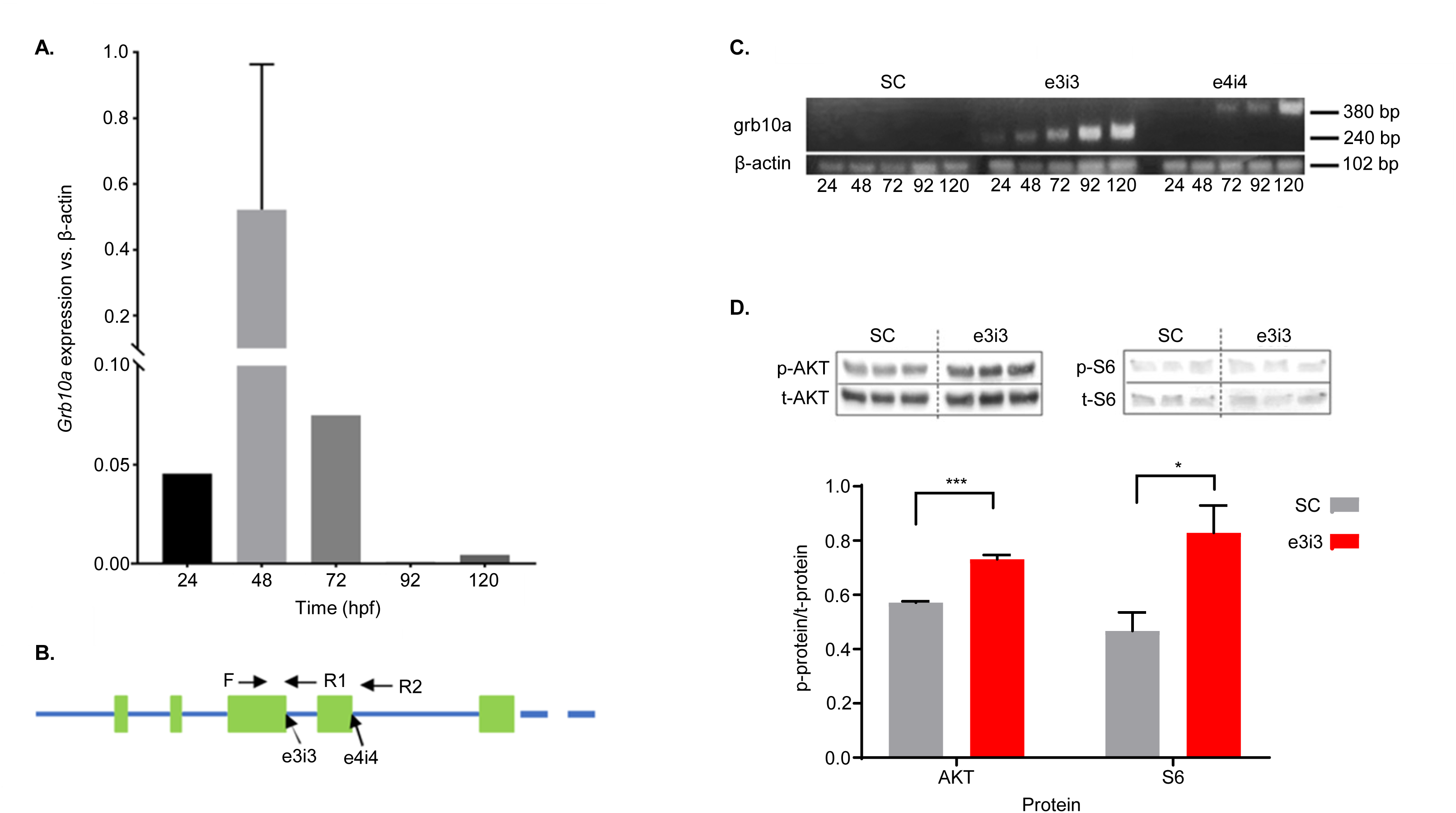
Grb10a is successfully knocked down in zebrafish injected with splice-blocking antisense oligonucleotides. ***1a*.** *Grb10a* qPCR of WT embryos (24-120 hpf, triplicated, n = 5 embryos per well). Data is shown as gene expression relative to β-actin and shows a significant peak at 48 hpf. ***1b.*** Schematic of the first five exons of the zebrafish *grb10a* gene. 5’ splice sites are highlighted with the forward and reverse primer triad indicated. ***1c.*** Multiplexed PCR amplification of the e3i3 and e4i4 splice site in embryos treated with either Standard Control morpholino, e3i3, or e4i4. β-actin was used as a positive control. ***1d.*** Western blot results of phosphorylated vs total protein ratios for two major signalling molecules of the insulin signalling pathway: AKT and S6. Quantitation using densitometry depicts mean + SEM. Activation of both proteins was found to be significantly elevated in KD zebrafish compared to SC (n = 3, unpaired t-test *** p = 0.0007, * p = 0.0413).

To knock down *grb10a* expression, zygotes were microinjected with splice-blocking antisense oligonucleotides e3i3 and e4i4. Exon 3 and 4 donor splice sites (*Figure 1b*) were targeted in order to confirm the specificity of the phenotype, in accordance with current guidelines^21^. Multiplexed RT-PCR amplification using primers flanking the splice sites (*Figure 1b*) showed a single product of the anticipated size for e3i3 and e4i4 embryos, and no product was detected for SC embryos (*Figure 1c*), consistent with successful incorporation of the corresponding intron.

To confirm grb10a KD induced a quantifiable impact on the downstream insulin signalling pathway, phosphorylation of key proteins was analysed by Western Blot. As shown in *Figure 1d*, phosphorylated (active) versus total protein ratios of AKT and S6 were significantly elevated in *grb10a* KD zebrafish at 96 hpf compared to SC (p = 0.0007 and 0.0413 respectively, n=3), consistent with the expected impact of grb10a KD.

### Growth Trajectory and Early Life Cardiometabolic Phenotype is Significantly Impacted by Transient *grb10a* Perturbation

To determine the effect of *grb10a* KD on growth, total body length was measured at 24- hour intervals over the first 5 dpf. As shown in *Figure 2a,* total body length was initially comparable between KD and SC zebrafish (2.857 ± 0.0549 mm vs 2.826 ± 0.0962 mm, p=0.7896, n=9 and n=10 respectively). Subsequently, KD zebrafish began to diverge from the SCs at 48 hpf, corresponding to the peak in *grb10a* expression observed in WT zebrafish (*Figure 1a*). KD zebrafish were longer on average than SCs (3.411 ± 0.0165 mm vs 3.177 ± 0.0231 mm at 72 hpf, p<0.0001, n=46 and n=41 respectively). This phenotype was reversed in zebrafish overexpressing *grb10a,* which were significantly shorter than SC counterparts (3.361 ± 0.0239 mm vs 3.505 ± 0.0339, p=0.001, n=24 and n=25 respectively), as shown in *Figure 2b*. Co-injection of e3i3 and *grb10a* RNA returned body length to SC levels (3.505 ± 0.0339 mm vs 3.564 ± 0.0265 mm, p=0.3792, n=25), confirming the validity of e3i3 induced *grb10a* KD. Moreover, *grb10a* KD induced by e4i4 (*Figure 2c*) also resulted in increased body length, and the ability of *grb10a* overexpression to suppress growth was shown to be dose dependent (*Figure 2c*). Intriguingly, by 120 hpf, body length converged, indicating activation of compensation to regulate growth and return to an “ideal” length post-hatch.

**Figure 2:**
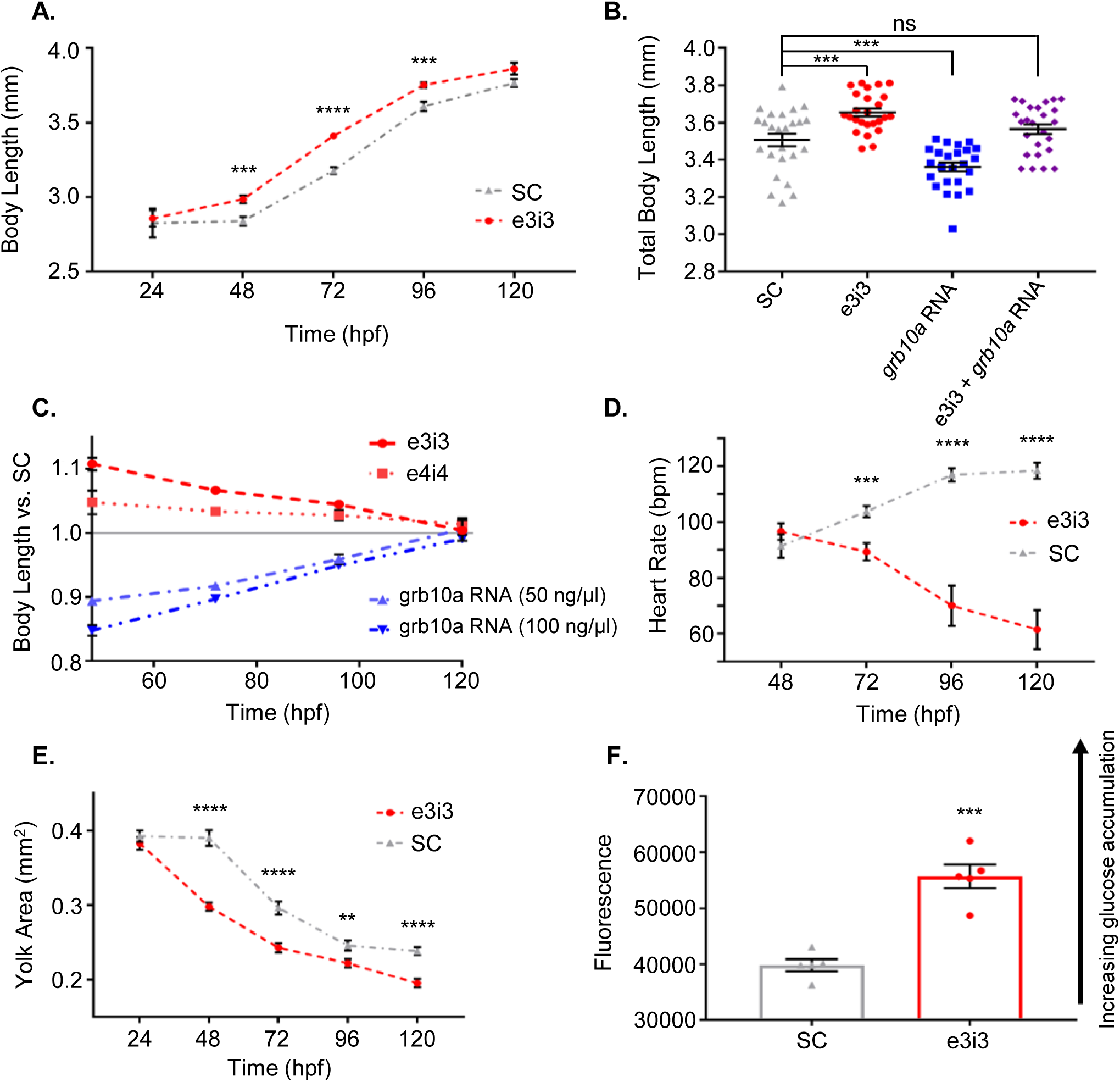
Growth and cardiometabolic phenotype are significantly impacted by grb10a perturbation. ***2a.*** Mean total body length ± SEM of SC and KD zebrafish from 24 to 120 hpf. T-test significance *** = 0.0006 **** < 0.0001 *** = 0.0001. ***2b*.** Mean total body length ± SEM and individual data points of 96 hpf zebrafish embryos (n = 25). *Grb10a* KD phenotype was reversed in *grb10a* overexpression zebrafish. Coinjection resulted in phenotype rescue. One-way ANOVA revealed KD zebrafish were significantly longer, while *grb10a* overexpression zebrafish were significantly smaller than SC (*** = 0.0001). Rescue zebrafish were of similar length to SC (ns = 0.3792). ***2c.*** Mean body length measurements relative to SC ± SEM. E3i3 and e4i4 both exhibit a propensity to elevate body length, while *grb10a* RNA shows a dose-dependent ability to inhibit body length. ***2d.*** Mean heart rate ± SEM in beats per minute of SC and KD zebrafish. Heart rate was significantly lower in KD embryos compared with SC after 48 hpf (*** = 0.0006, **** < 0.0001). ***2e*.** Mean yolk area ± SEM of SC and KD embryos over the embryonic life stage. Following the initial 24 hours, KD zebrafish had significantly smaller yolks compared to SC, indicating yolk content was metabolised at a much higher rate (** = 0.0084 **** < 0.0001). ***2f*.** Glucose Uptake-Glo^TM^ Assay of 96 hpf KD and SC zebrafish, where higher luminescence indicates a greater accumulation of intracellular 2D6P. Luminescence was approximately 30% greater in KD zebrafish compared to SC (*** = 0.0002). All SC vs KD comparisons by unpaired t-test.

To investigate the impact of *grb10a* KD on the developing cardiac system, heart rate was measured over the first 5 dpf. As shown in *Figure 2d*, average heart rate began to diverge between the treatment groups at 48 hpf, again correlating with the WT peak in *grb10a* expression. While heart rate increased slightly over time in SC zebrafish, in-line with increasing body size, average KD heart rate fell. By 120 hpf, average heart rate was almost 50% lower in the KD compared to SC (61.5 ± 6.97 bpm vs 118.4 ± 2.83 bpm, p<0.0001, n=14 and n=19 respectively).

To determine whether *grb10a* KD had an impact on metabolic rate, yolk absorption was measured to indicate energy demand. As shown in *Figure 2e*, there was initially no difference in yolk area between the groups (p=0.8185, n=10). As with body length and heart rate, SC and KD yolk consumption began to diverge at 48 hpf (0.298 ± 0.0055 mm^2^ vs 0.390

± 0.0104 mm^2^, p<0.0001, n=18 and n=19 respectively). The elevation in yolk consumption observed in the KD fish suggests an elevated metabolic rate. To support this conclusion, a Glucose Uptake-Glo^TM^ Assay was performed to compare the rate of glucose uptake. As shown in *Figure 2f*, 2D6P accumulation was significantly higher in KD zebrafish compared to SC, an increase of almost 30% (p=0.0002, n=5), indicating glucose uptake was elevated. These findings are consistent with the role of *grb10a* as a negative regulator of the insulin signalling pathway^55^ and a coordinator of growth and metabolism.

### Transient *grb10a* Knockdown Persistently Dysregulates Age-Associated Gene Expression

To understand whether the observed changes were coupled with lasting changes in gene expression, the transcriptomic landscape of SC and KD zebrafish was investigated over the first 30 dpf. Unsupervised hierarchical clustering, standard deviation filtering, and maximised projection scores (MPS) were used to define a set of genes with strong age-association in the SC zebrafish (297 genes, MPS = 0.43). These genes fell into four distinct clusters, associating with 5, 10, 15, and 20-30 dpf (163, 15, 32, and 87 genes respectively) (*Figure 3a*).

**Figure 3.**
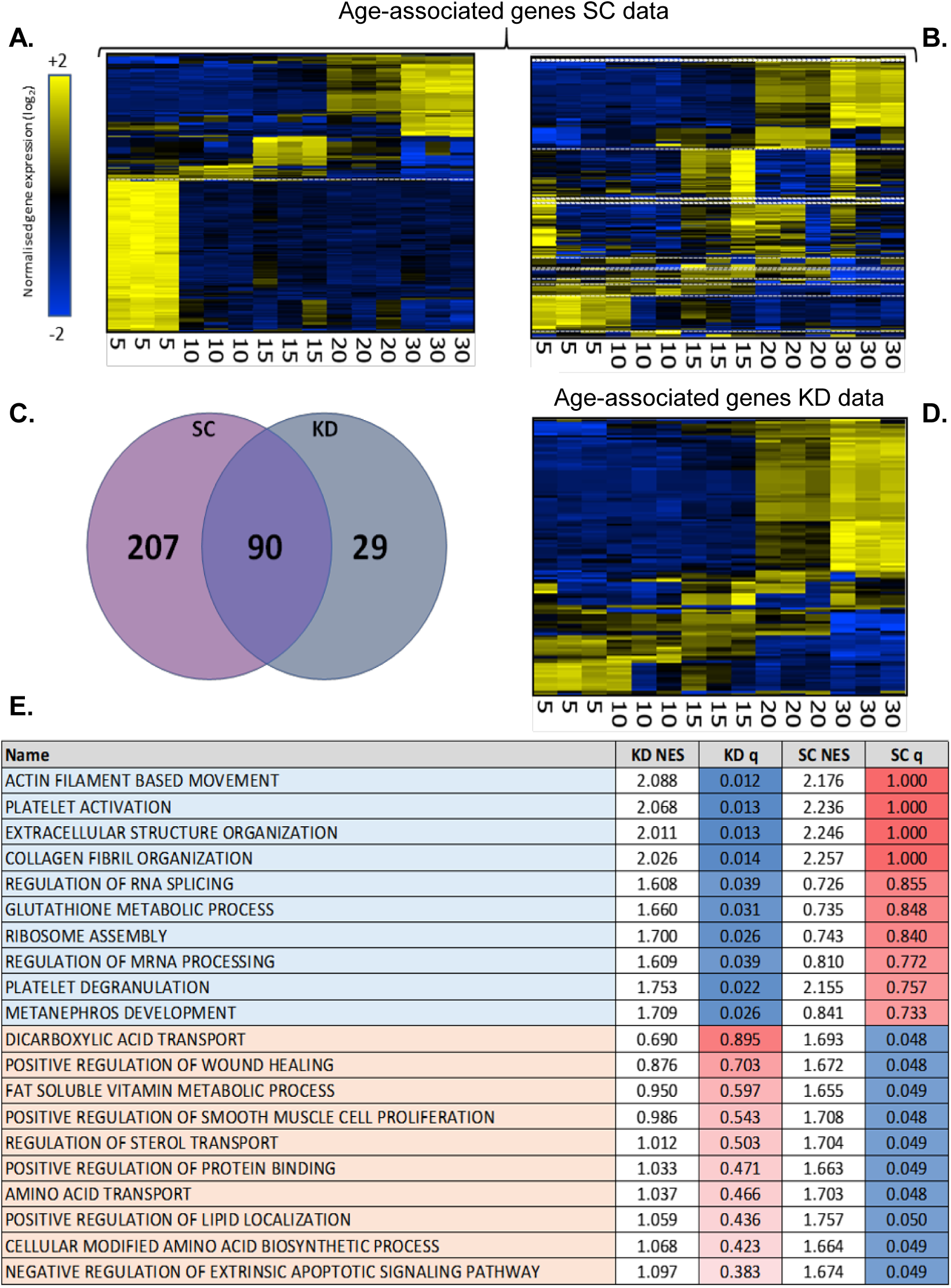
Transcriptomic analysis of Standard Control and grb10a Knockdown gene expression over the first 30 dpf. Hierarchically clustered heat maps of gene expression generated from an Affymetrix GeneChip™ Zebrafish Genome Array of SC (***3a***) and KD (***3b*** and ***3c***) zebrafish RNA, taken at 5, 10, 15, 20, and 30 dpf. Expression segregates into three clusters in SC zebrafish (***3a***). Clustering of the same genes identified in ***3a*** are disrupted in the KD dataset (***3b***). Analysing the KD dataset independently shows age-related hierarchical gene expression in the falls into two clusters (***3c***). ***3d***. Venn diagram of age associated genes. ***3e***. Gene set enrichment analysis, using the GO Biological Process Ontology gene list, of the age-related genes in the SC and KD datasets. The top 20 most enriched pathways with differential expression are included here, with associated normalised enrichment scores and q-values.

When performing the same analysis in KD zebrafish, the 5 dpf cluster was significantly disrupted (*Figure 3b*). While the clustering was largely conserved in the latter three clusters (10 dpf – 14/15, 15 dpf - 29/32, and 20-30 dpf – 74/87 genes), genes strongly associated with 5 dpf in the SCs were generally expressed at different time points in the KDs (dotted white lines, *Figure 3a* and *Figure 3b*), with a mapping of only 38/163 genes (23%). Notably, five genes associated with 5 dpf in SC zebrafish mapped to 20-30 dpf in the KD. This suggests the expression of these genes is usually associated with early larval development, but instead, in the KDs, is associated with the late-juvenile stage. Human orthologues of these dysregulated genes include *DGAT2* (fatty acid metabolism), *GAMT* (energy storage, muscle contraction, and fatty acid oxidation), and *PDIA2* (thiol-disulphide interchange, particularly in the pancreas).

As the SC gene clusters were disrupted in the KD dataset, unsupervised analysis of the KD data was performed to identify the subset of age-associated genes in the KD zebrafish. 119 genes were identified (MPS = 0.37) which segregated into 5-15, 15-30, and 20-30 dpf (37, 16, and 66 genes) (*Figure 3c*).

To assign functionality to these age-related genes, Gene Set Enrichment Analysis was performed for genes identified in both the SC and KD datasets. Functionality conserved between the SC and KD is described in *Supplemental Table 1*. Dissimilar pathways (*Figure 3e*) included several actin and collagen related pathways, and extracellular structure and RNA processing, which were age-related in KD but not SC zebrafish. Conversely, several metabolic pathways were age-associated in SC zebrafish but not in KD animals. This clear dysregulation of age-related gene expression in the KD zebrafish implies transient *grb10a* KD induces persistent remodelling of the transcriptome.

### Transient *grb10a* Knockdown Disrupts Transition Between Larval Gene Clusters

To investigate the co-ordination of the whole transcriptome with age-associated gene expression, hypernetwork models were constructed for the SC and KD datasets based on the gene sets identified in the cluster analysis (297 and 119 genes respectively). This approach has been used to model functional relationships^31^. *Figure 4* describes the results, where colour intensity represents the number of shared interactions between each gene pair. In SCs, shared interactions segregated into three groups, correlating to 5, 10-15, and 20-30 dpf (161/297, 50/297, and 86/297 genes) (*Figure 4a*). In KD animals, however, genes segregated into only two groups, either 5-15 dpf or 20-30 dpf (31/119 and 65/119 genes) (*Figure 4b*). Notably, the large group of co-ordinated interactions at 5 dpf was absent in the KD dataset, with a combined group of genes correlating with 5-15 dpf identified instead.

**Figure 4:**
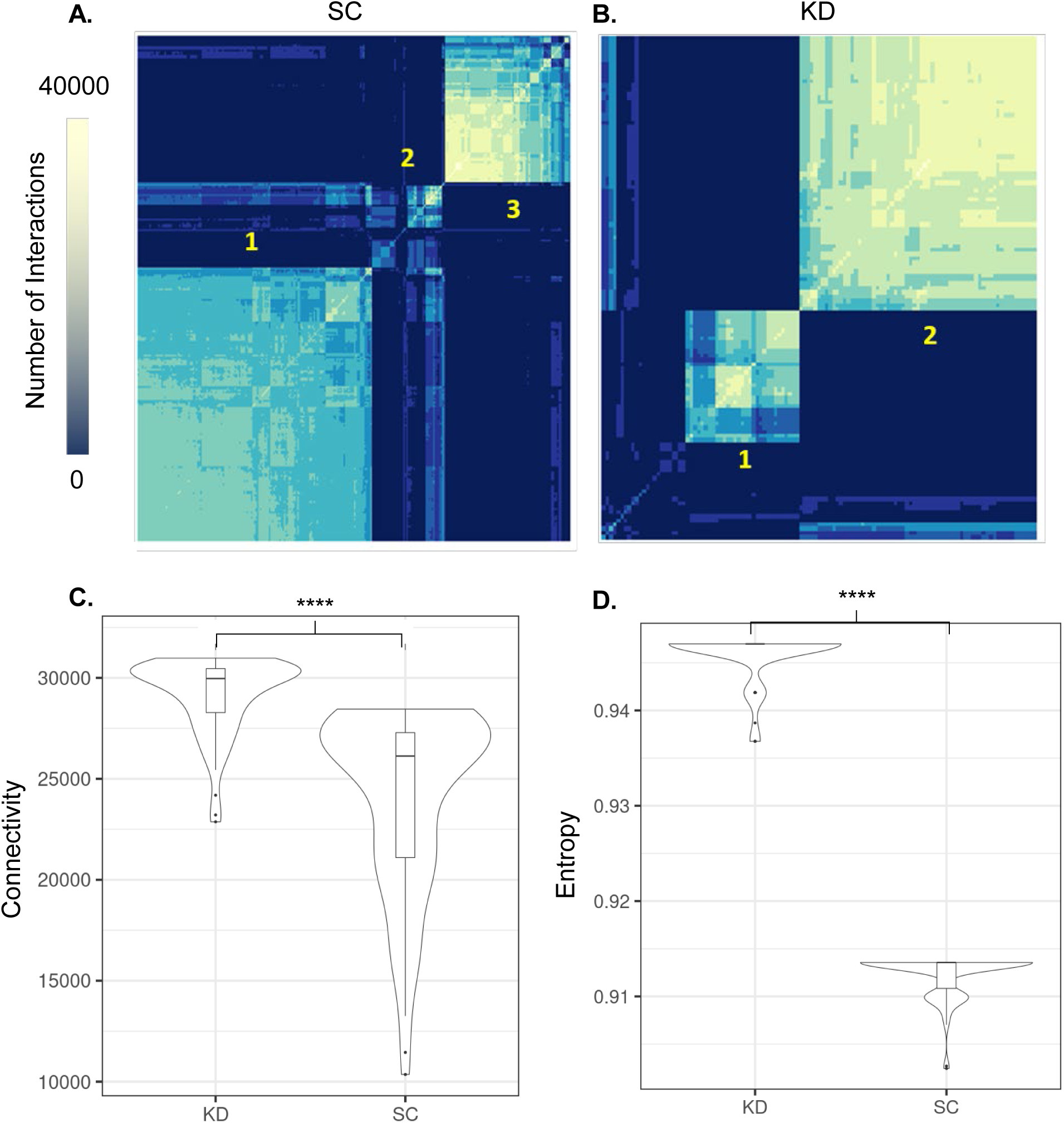
Coordination within the transcriptome is altered by grb10a KD, as assessed by hypernetwork analysis. **4a.** Hypernetwork analysis defined three clusters of highly connected genes in Standard Control zebrafish, corresponding to 5 dpf, 10-15 dpf, and 20- 30 dpf respectively. ***4b***. Two clusters were defined in the KD data, corresponding to 5-15 dpf and 20-30 dpf. ***4c*** *and **4d**.* Violin plots of connectivity (left) and entropy (right) in the 20-30 dpf cluster. The KD transcriptome was more connected with greater entropy compared to Standard Controls, suggesting more diverse functionality.

### *Grb10a* Knockdown is Associated with Increased Connectivity and Crosstalk Within the Transcriptome

To quantify these differences in the co-ordination of the transcriptome, two network topology parameters were used: connectivity and entropy. Hypernetwork connectivity quantifies the number of higher order interactions within the transcriptome and is related to function^31, 56^, while entropy is a measure of information content and serves as an indicator of “disorder”. I.e. lower entropy (more order) describes a network with little crosstalk and a more discrete function^34, 57^, while a network with greater entropy has increased crosstalk and pleiotropic functions.

Changes in connectivity (*Figure 4c*) and entropy (*Figure 4d*) were calculated for genes identified as active at 20-30 dpf by the previously described analyses. The impact of KD on the co-ordination of age-associated genes in the network was identified by comparing the connectivity and entropy of SC and KD datasets. To provide a comparison between age-associated genes and genes associated with other functions, connectivity and entropy were also calculated for randomly selected gene sets (*Supplemental Figure 2*).

Genes associated with 20-30 dpf were more highly connected and less entropic (more ordered) than random genes in both the KD and SC datasets (p < 2.2 x10^-16^). Comparison of age associated genes in KD and SC revealed a higher entropy (1.05-fold, p < 2.2 x10^-16^) and higher connectivity in the KD (1.20-fold, p < 2.2 x10^-16^). This suggests age-associated genes in the KD zebrafish transcriptome have more interactions and are less ordered (have more crosstalk) than in the SC.

### Pathways Associated with Transcriptome-Wide Remodelling Support an Alteration in Cardiometabolic Phenotype

Having identified a core set of age-related genes, a broader set of genes which coordinate with these genes was defined as the complete subgraph between the central age-related genes of the hypernetwork and their correlates. 12775 (KD) and 459 (SC) genes were defined in this set from the 20 to 30 dpf hypernetwork clusters described previously, all significantly associated with age (rank-regression: KD q < 1.50 x10^-6^, SC q < 4.44 x10^-2^; *Supplementary Table 2*). 28 times more genes were implicated in the KD. Both datasets showed a skew towards a positive association with age (68% SC vs 59% KD) (*Figure 5a, Figure 5b*). The overlap in gene expression between the two datasets was 244 (77.9% of SC) and 60 (41.1% of SC) in the positively and negatively correlated sets, respectively. Thus, genes in the wider transcriptome with co-ordinated expression at 20-30 dpf in SC zebrafish demonstrated a similar pattern of expression in KD (*Figure 5c*). However, the inverse is not true, as genes with co-ordinated expression at 20-30 dpf in the KD zebrafish were dysregulated in the SC (*Figure 5d*).

**Figure 5:**
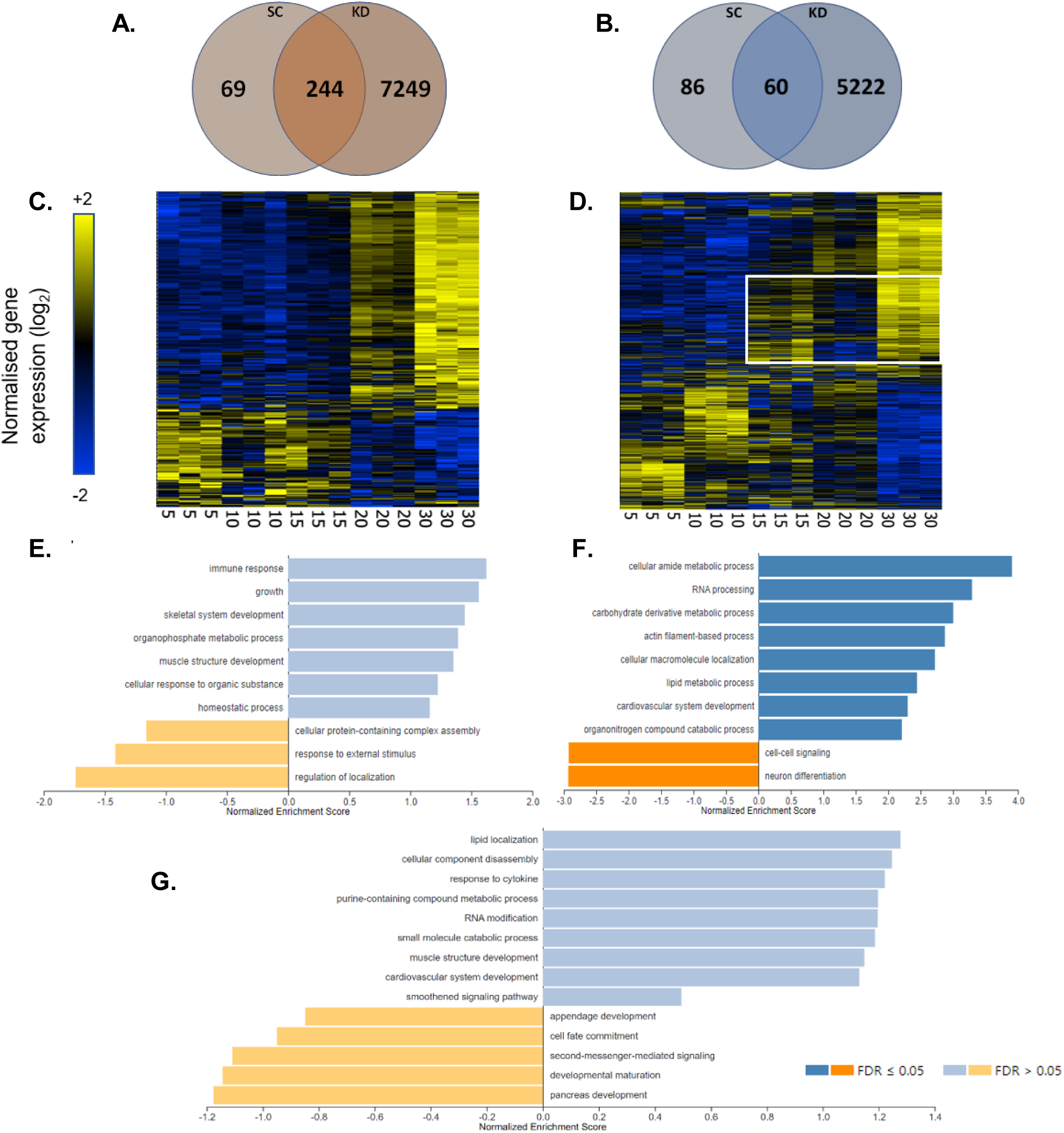
Analysis of the set of genes in the wider transcriptome shows a 27.8-fold increase in the KD ZF. ***5a-5b***. Venn diagrams of genes positively (***a***) and negatively (***b***) correlating with age. ***5c-5d***. Hierarchically clustered heat maps of gene expression of the genes identified in the wider transcriptome. Gene expression in the standard control (***5c***) cluster into two age related groups, whereas expression in the knockdown (***5d***) show significant dysregulation. ***5e-5f*** Gene set enrichment analysis (GSEA) ranked by R-value of rank age regression in the standard control (***5e***) and knockdown (***5f***). ***5g*** GSEA ranked by R-value of rank age regression of the cluster of genes in the white box in ***5d***.

To assign functionality, GSEA was performed as previously described (ranked by R-value, top 100 pathways with weighted set cover). Results of this ontology analysis are outlined in *Figure 5e* and *Figure 5f*. The two datasets featured distinctly different regulated pathways. RNA processing, a variety of metabolic pathways, and cardiovascular development featured in the KD dataset, whereas growth and immune signalling were featured in the SC dataset. Functional analysis of these determined that the greatest differences in activity between the KD and SC were in *Positive Regulation of Lipid Localization*, *Amino Acid Transport* and *Cellular Modified Amino Acid Biosynthetic Process* (*Figure 6*).

**Figure 6:**
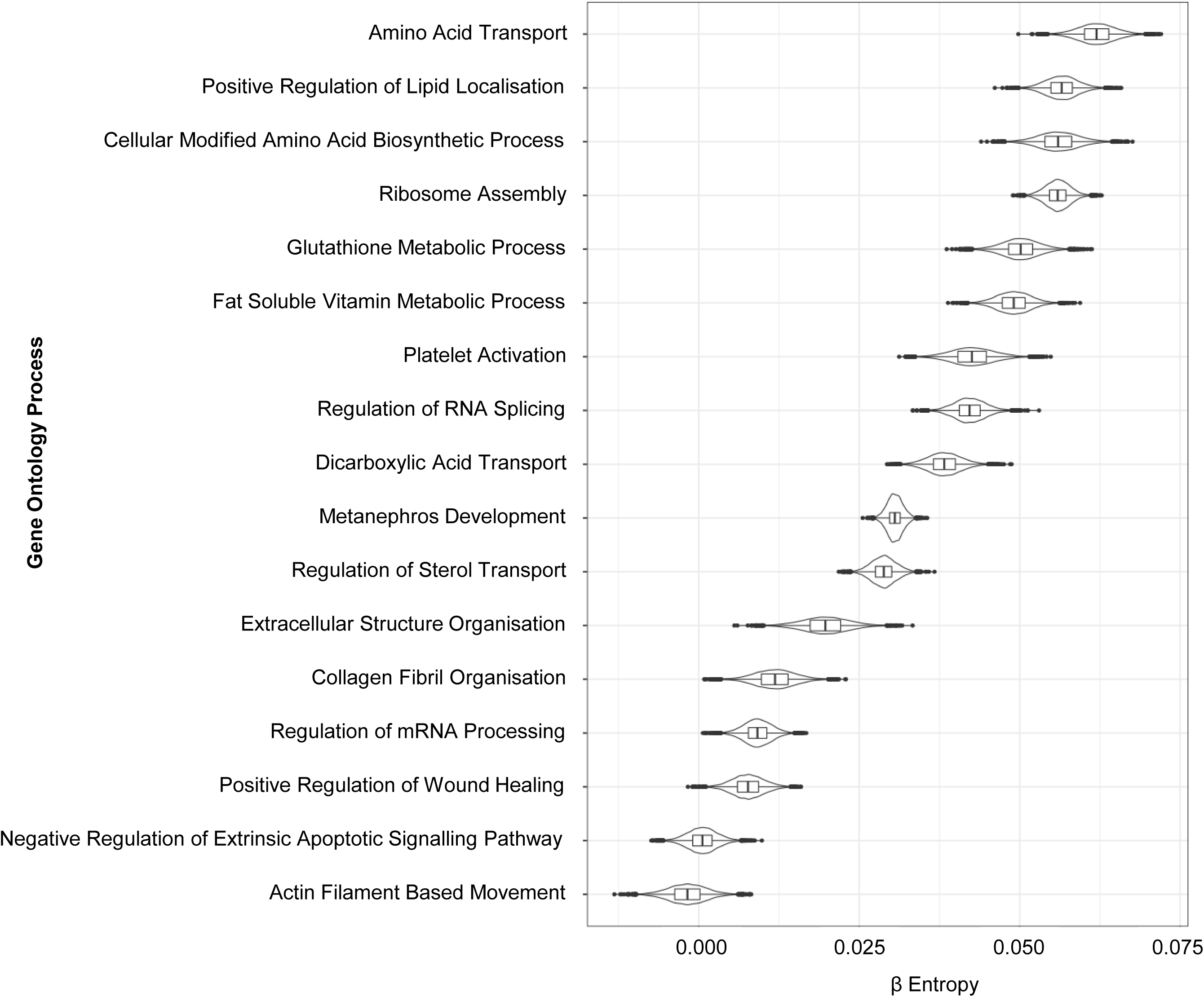
Hypernetwork modelling of ontology highlights pathways with functional difference between SC and KD. Hypernetworks were iteratively generated from genes attributed to each pathway and entropy was modelled across the two treatment groups. β values represent the difference in entropy between SC and KD, with a β value of 0 indicating no difference between groups. Ontology classes are considered to be significantly different if the distribution of β values (89% CI) does not include 0. *Actin filament based movement* and *negative regulation of extrinsic apoptotic signalling pathway* were the only two pathways with no functional difference identified between the groups.

A distinct pattern of gene expression was identified in the group of genes dysregulated at 20-30 dpf in the KD dataset (white box, *Figure 5d*). These genes were upregulated at 15 dpf, downregulated at 20 dpf, and re-upregulated at 30 dpf. This subgroup of 3460 genes (*Supplemental Table 3*) was associated with a range of gene ontologies related to metabolism and development (*Figure 5g*) and was synchronous with the spike in growth identified in *Figure 7a*. Specific pathways following this pattern of expression included cardiovascular system development, muscle structure development, and developmental maturation.

**Figure 7:**
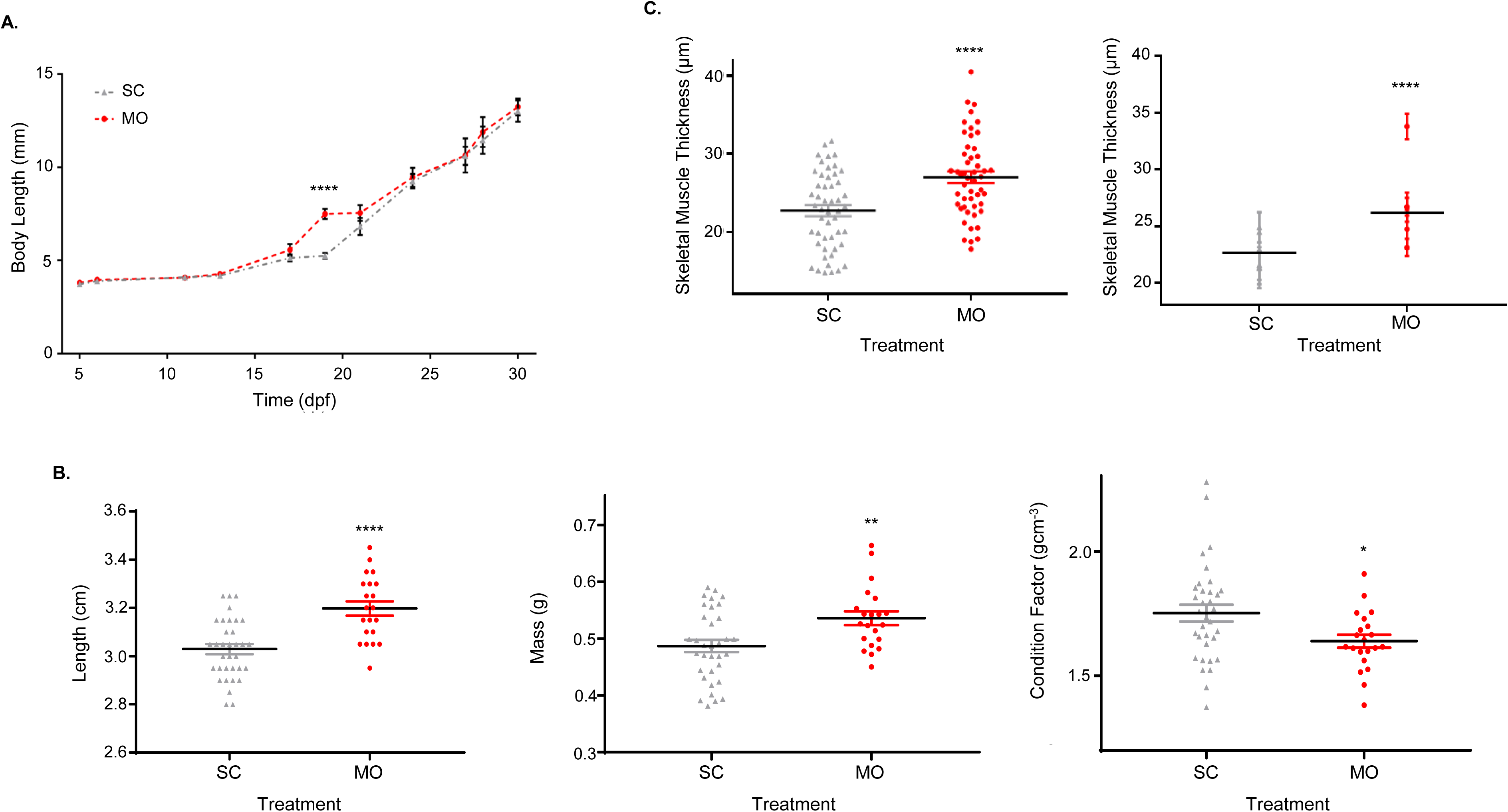
Length and cardiac function are permanently altered. ***7a.*** Mean total body length of SC and KD zebrafish up to 30 dpf. Following the embryonic growth spurt, KD zebrafish experienced an additional period of rapid growth between 15 and 20 dpf (**** < 0.0001). ***7b.*** Individual total body length (left), mass (middle), and condition factor scores (right) for 18-month KD and SC zebrafish (n = 21-24). Length and mass were significantly higher in the KD (**** p < 0.0001, ** p = 0.005), while condition factor was significantly lower (* p = 0.0215), indicating KD zebrafish have leaner bodies. ***7c.*** Thickness of 10 individual skeletal muscle fibres stained with Masson’s Trichrome from 5 individuals (left) and means grouped by individual (right) taken from a position posterior to the dorsal fin. Fibres were consistently thicker in the KD zebrafish (**** p < 0.0001). All comparisons by unpaired t-test. Data are presented as mean or individual values ± SEM.

### The Impact of Transient *grb10a* KD Persists into Adulthood

As *grb10a* KD significantly impacted early-life growth, metabolism, and cardiovascular development and was associated with persistent remodelling of the transcriptome, investigation was conducted into the phenotypical differences in older zebrafish.

Total body length measurements up to 30 dpf are shown in *Figure 7a*. Growth was significantly elevated in the KDs compared to SCs between 15 and 20 dpf (30% increase, 7.493 ± 0.2726 mm vs 4.320 ± 0.1594 mm, p < 0.0001, n = 10). This corresponded to the cluster of dysregulated genes identified by hypernetwork modelling (*Figure 5g and white box in Figure 5d*). The growth profile of the KDs was shifted to an earlier age compared to the SC, suggesting a faster rate of maturation. The GSEA highlighted a reduction in activity in developmental maturation pathways (normalised enrichment score < 1) in SCs vs. KDs, which was reflected in the growth rate at approximately 20 dpf.

Late-life body length and mass measurements were recorded in 18-month zebrafish to investigate the lasting impact of *grb10a* KD on the phenotype. Final body length (*Figure 7b.*) was higher, with KDs 1.7 mm longer than SCs (3.20 ± 0.029 cm vs 3.03 ± 0.021 cm, p < 0.0001). These fish were also approximately 10% heavier (0.54 ± 0.012 mg vs 0.49 ± 0.011 mg, p = 0.005). On the other hand, the Fulton’s condition factor (an indicator of body condition) was closer to 1 in the KDs (1.64 ± 0.026 vs 1.75 ± 0.034, p=0.0215) despite the increase in mass. To equate this to differences in body composition, skeletal muscle fibres, isolated from the base of the dorsal fin, were embedded and stained with Masson’s Trichrome (*Figure 7c.*). The average muscle fibre diameter was approximately 20% greater in the KDs (27.01 µm ± 0.728 vs 22.71 µm ± 0.698, p < 0.0001), likely contributing to the increase in mass despite a decrease in roundness.

The impact on cardiac health was assessed by qPCR and histology. As shown in *Figure 8a*, *myl7* expression (an index of muscle mass and hypertrophy) in the heart was over 20% greater in KDs (p < 0.0001, n = 3), while *nppa* expression (activated in response to ventricular stress during hypertrophy and heart failure^58–60^) was reduced by approximately 40% (p = 0.0012, n = 3). There was no difference in *pcna* expression (proliferating cell nuclear antigen), suggesting there was no difference in proliferation in the cardiac tissue. To support these findings, ventricular morphology was also assessed. As shown in *Figure 8b*, the ratio of compact myocardial layer to trabeculated was significantly greater in the KD zebrafish (0.369 +/- 0.07 vs 0.185 +/- 0.03, p-value = 0.0288). This was coupled with an overall increase in tissue density (76.3% vs 64.4%, p-value = 0.0289).

**Figure 8:**
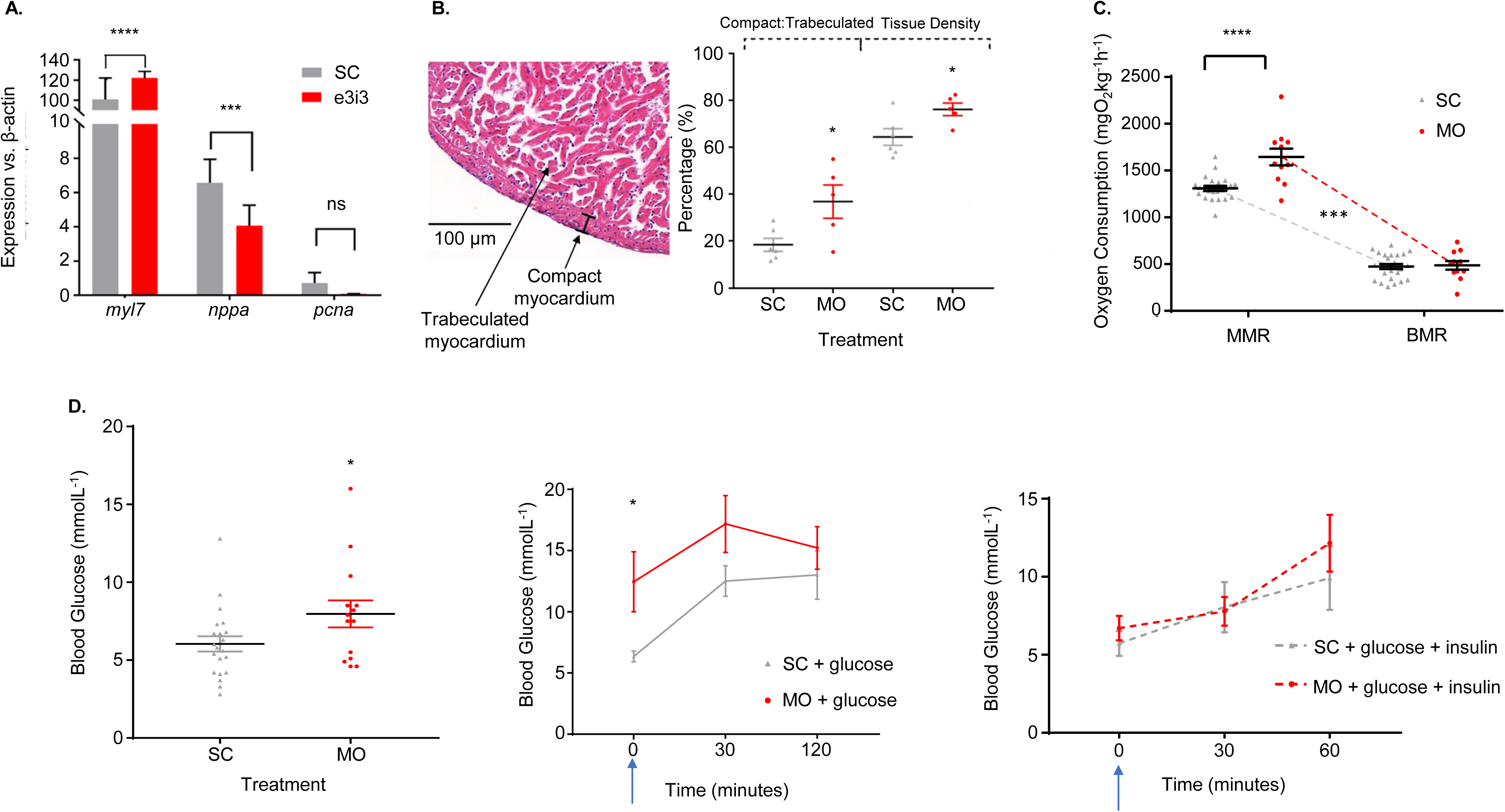
Changes established during embryonic development perpetuate into variations in adult function. ***8a.*** qPCR results of three genes associated with cardiac performance in adult cardiac tissue, relative to β-actin. *Myl7* expression was significantly elevated in KD zebrafish (**** p < 0.0001) while *nppa* expression was significantly down regulated (*** p = 0.0012) compared to SC zebrafish. There was no significant difference between the expression of *pcna* (p = 0.3041). 8***b.*** Ventricular morphometrics obtained by Masson’s Trichrome histology comparing KD and SC compacta thickness and tissue density. The compacta layer was significantly thicker (* p = 0.0288) and overall density was higher (* p = 0.0289) in the KD zebrafish, suggesting greater cardiac efficiency (n = 5-6). ***8c***. Maximum (MMR) and Basal (BMR) oxygen consumption of adult (18 month) zebrafish, adjusted for body mass. BMR was comparable between the two groups, while MMR was greater in the KD zebrafish (**** p < 0.0001), resulting in a greater aerobic scope (dotted line, *** p = 0.0007). **8*d.*** Fasting blood glucose concentrations of adult zebrafish from each treatment group (n = 14-21). KD zebrafish had significantly higher circulating blood glucose levels (* p = 0.0463). ***8d.*** Glucose tolerance (top) (n = 10) and insulin sensitivity (bottom) (n = 8–12) trials in adult (18 month) KD and SC zebrafish. The treatment groups were equally able to tolerate glucose challenge and responded similarly to insulin. Treatment started immediately after the first measurement. Data are presented as either mean or individual values ± SEM, and all comparisons are by unpaired t-test.

To determine whether metabolic rate was persistently altered, stop-flow respirometry was conducted on juvenile (30 dpf). As shown in *Figure 8b*, KDs consumed 7 times more oxygen than SCs (134.4 ± 15.60 µgO_2_h^-1^g^-1^ vs 19.0 ± 3.28 µgO_2_h^-1^g^-1^, p < 0.0001, n = 5), suggesting metabolic rate remained significantly elevated in juvenile fish.

To determine whether this was a permanent change, adult (18-month) zebrafish were also investigated. Peak and basal oxygen consumption were recorded to further investigate the “lean” phenotype. Maximum oxygen consumption, achieved following exhaustive activity, was elevated by ∼25% in KDs, as shown in *Figure 8c* (1645 ± 90.11 mgO_2_kg^-1^h^-1^ vs 1309 ± 26.48 mgO_2_kg^-1^h^-1^, vs p < 0.0001). There was no significant difference in basal metabolic rate (486.5 ± 46.75 mgO_2_kg^-1^h^-1^ vs 474.1 ± 27.78 mgO_2_kg^-1^h^-1^, p = 0.8132). Consequently, aerobic scope was greater in the KDs (1158 ± 100.7 vs 810.1 ± 42.95, p = 0.0007), supporting the conclusion that the fish conform to a “leaner” phenotype.

As grb10a is involved in regulating the insulin signalling pathway, glucose tolerance and insulin sensitivity tests were performed on adult zebrafish (18 months) to determine whether there was a lasting biological impact. Fasting blood glucose (*Figure 8d*) was significantly higher in KDs (7.96 ± 0.867 mmolL-1 vs 6.04 ± 0.493 mmolL-1, p = 0.0463). Glucose tolerance testing showed both groups produced a similar response to glucose challenge and were similarly able to respond to introduction of insulin.

## Discussion

The first key finding from this study is that grb10a regulates embryonic growth in zebrafish, consistent with its role in mammalian embryogenesis^47–49^, and transient knockdown is sufficient to have an impact on downstream pathways. This is consistent with its role as a negative regulator of insulin signalling^50–51^. Western Blotting showed a clear upregulation in insulin signalling pathway activation downstream of grb10a, which was coupled with significant changes to the early phenotype, including elevated growth rate, increased metabolism, and distinct cardiac alterations.

The observed changes in phenotype coincided with the peak in grb10a expression, confirming the role of *grb10a* as a coordinator of embryonic growth and development. The elevated growth coincided with higher metabolic rate, consistent with an increase in energy demand due to a larger population of highly proliferating cells. Cardiac changes were also present, supporting the conclusion that grb10a has a role in coordinating growth, metabolism, and cardiac development. This is the first study, to our knowledge, to provide *in vivo* evidence for the coordination of these distinct pathways.

This study also provides clear evidence for a compensatory growth mechanism activated during early larval development, which regulates adherence to an “ideal” body length. Both elevated embryonic growth and growth suppression were compensated for by 5 dpf, with body length returning to comparable values in all treatment groups. This suggests there may be a benefit to entering life at this controlled size. The altered phenotype established by day 5 served as a basis for life-long changes to the organism. Body length, metabolism, and cardiac physiology were all significantly altered in 1.5-year-old zebrafish, despite no further manipulation to the organism. This provides essential, novel, *in vivo* evidence for the Fetal Origins of Adult Health hypothesis, which proposes that events during development can have life-long impacts on an organism.

Furthermore, this study has shown for the first time that a transient disruption in the expression of a single gene can result in permanent remodelling of the transcriptome. This highlights the importance of regulated control during embryogenesis and the significant impact small changes, such as increase in growth, can have on the developed organism. Age-associated genes in both transcriptomes showed increased connectivity and decreased entropy compared to non-age associated genes. This demonstrates that the genes identified as age-associated are better connected to one another and more ordered in their interactions than randomly selected genes. Entropy and connectivity were both higher in KD than SC which demonstrates that age-associated genes share more interactions in KD but with less discrete organization than in SC. These results support the conclusion that the KD transcriptome is dysregulated, compared to the SC transcriptome.

Notably, gene expression at 5 dpf was significantly dysregulated in the KD zebrafish compared to the SCs. An early transition in gene expression identified in the SCs, occurring between 5 and 10 dpf, likely corresponding to a shift away from early developmental pathways, was lost in KD zebrafish. This loss was reflected both in the age-related genes and the coordination of the wider gene set. This fundamental difference in the transcriptome during larval development was particularly notable in a subset of genes with fluctuating gene expression in the KD zebrafish. Genes associated with cardiovascular development, muscle development, and developmental maturation showed a distinct pattern of upregulation at 15 dpf, downregulation at 20 dpf, and re-upregulation at 30 dpf, coinciding with the spike in growth in the larval zebrafish. This pattern may contribute to the left-shift in growth rate observed in the KD zebrafish, which is reflected in human cohorts experiencing early puberty^61^. Furthermore, functional differences in cardiac and skeletal muscle and metabolic phenotype were confirmed to persist into the adult zebrafish, suggesting these transient changes in gene expression have long-lasting implications for the organism.

This model of embryonic growth perturbation may also yield significant insights into the propensity for early growth disruption to correlate with increased risk of cardiovascular and metabolic disease in later life^62–65^. It is widely accepted that many disorders are likely to have their origins during embryonic development^66–69^, but the mechanisms involved are not fully understood, and targeted *in vivo* research is lacking. Mammalian models have been used to investigate the immediate impact of embryonic growth disruption^69–71^, findings which are replicated in this study. However, longitudinal studies are absent from the literature, and little research has been conducted into the whole-life significance of early growth disruption. Multiple distinct growth trajectories can yield similar birth weights, including catch-up and catch-down growth^62^ (both of which were achievable by modulation of grb10a expression). Catch-up and catch-down growth have been reported to correlate with increased risk of chronic health disorders^65, 72, 73^, and as *grb10a* modulation is sufficient to alter embryonic growth trajectory, metabolic rate, and cardiac function, the model generated in this study may prove key in understanding the mechanisms involved in the developmental origins of health and disease and identifying novel avenues for prevention and treatment.

As identified in mammalian studies^19, 20^, *grb10a* KD was associated with a “leaner” phenotype. KD zebrafish displayed an elevation in skeletal muscle fibre thickness and an increase in body mass, likely as a result of altered body composition in favour of skeletal muscle. Body condition also conformed to a leaner morphology in the KDs, where Fulton’s condition factor, used by fisheries and in research to assess the condition of fish stocks^74^, was closer to 1. Healthy fish in good condition have condition scores close to 1, while higher or lower scores indicate overly fat or “skinny” fish respectively. Together, these findings support the existence of a “lean” phenotype associated with suppressed grb10a activity and greater insulin signalling activation. This is expected, as circulating insulin levels positively correlate with growth, and serve as an indicator of resource availability in the mature environment.

This study further extended the “lean” phenotype to include cardiovascular and metabolic changes, the first *in vivo* work to combine these aspects. The establishment of an altered cardiac phenotype during embryogenesis was followed by a notable change in ventricle morphology. To maintain optimal cardiac output, a reduction in the rate of contraction must be paired with increased cardiac efficiency and an increase in force or volume of contraction. The increased demand on the cardiac tissue resulting from the reduction in heart rate resulted in compacta thickening and an increase in ventricular density. This is consistent with increased stroke volume and greater cardiac efficiency due to an increase in contractility. Gene expression analysis of cardiac tissue, coupled with histological evidence, supported this conclusion, and indicated the increase in musculature was due to an increase in cell size rather than cell number. The greater degree of *myl7* expression in the absence of an increase in *pcna* expression suggests an increase in hypertrophy (muscle size) rather than hyperplasia (cell number) in the cardiac tissue.

Aerobic scope was also elevated as a result of grb10a KD, consistent with a “lean” phenotype. This greater aerobic scope suggests the potential energy for non-essential activities is elevated in KDs. As a result, these fish may have the capacity to outperform SC counterparts in other energy-heavy tasks, such as swimming and courtship. The difference in peak oxygen consumption between the groups suggests there is either increased oxygen demand or an increased ability to extract oxygen from the environment in KD zebrafish.

Glucose homeostasis control was also altered in KD zebrafish. Glucose uptake was elevated during embryogenesis and energy stores were depleted more rapidly, which was reflected in elevation of fasting blood glucose in later life. Taken together with the elevation in oxygen consumption observed in juvenile and adult zebrafish, this indicates greater metabolic rate. This suggests KD zebrafish, while having a comparable basal energy demand, have a greater demand for energy when performing non-essential activities. This may be due to higher contribution of skeletal muscle to body composition, as heavily respiring tissues require more energy when active than other tissues. KD zebrafish showed a greater ability to maintain glucose homeostasis compared to SCs, responding more rapidly to glucose challenge. As metabolic rate was greater in the KD zebrafish, this increase in the rate of glucose clearance may be a direct consequence of heavily respiring tissue removing glucose from the bloodstream at a higher rate.

Findings from this study may also provide important for the agri- and aquaculture industries. The promotion of embryonic growth induced by *grb10a* KD may be a key alternative to growth hormone treatment for improving meat yield and production efficiency, and larger juvenile fish with greater aerobic scope are likely to be more capable of overwintering^75, 76^, and thus improve fish stocks. This is particularly key, as the impact of climate change on fisheries is an increasing global concern^77–79^. The increase in skeletal muscle contribution to body composition also has relevance for industry, as lean products have greater value and marketability, particularly as current health trends are increasing demand for low-fat products.

## Conclusion

This study proves for the first time that transient knockdown of *grb10a* expression is sufficient to permanently alter growth trajectory, metabolic rate, and cardiac physiology. We have presented significant evidence to suggest grb10a plays a previously unidentified, fundamental role in the coordination of these distinct physiological pathways and could represent a promising target for enhancement of meat yield and quality in agri- and aqua-cultural. Furthermore, this study shows that altering the expression of a single gene during embryogenesis can remodel the entire transcriptome. We have shown that remodelling established during embryogenesis provides a basis on which the adult phenotype is formed, clearly demonstrating the lasting impact of early-life events on the whole life course of an organism. This provides the first longitudinal *in vivo* support for the Foetal Origins of Health hypothesis in *Danio rerio*. We have also expanded on the “lean” phenotype associated with grb10 KO to include a distinctly altered metabolic phenotype and improved ventricular efficiency. These life-long alterations may have ongoing implications for survival, and the model generated in this study, featuring easily measurable phenotypic characteristics, a short developmental window, and rapid generation time, will be indispensable for future research into the mechanisms underpinning the foetal origins of health.

## Acknowledgements

This work was funded by the Biotechnology and Biological Sciences Research Council at the University of Manchester, Manchester, UK, in combination with an unrestricted Merck research grant, Darmstadt, Germany. Preliminary and the transcriptomic work was partially funded by a University of Manchester Research Institute Pump Priming Fund entitled “The quantitative comparison of the Zebrafish as a model of human development in relation to paediatric medicine”, RMS 104699.

The authors would also like to acknowledge the following people: Dr. Andrew Badrock for his training, guidance, and donation of reagents, and Jack Broadbent and Joseph Whitehead for their preliminary work in quantifying growth and metabolism in this model.

## Data and Code Availability

Transcriptomic data is available from the Gene Expression Omnibus (GSE162474). R code is available online at: https://github.com/terencegarner/GRB10_KD_ZF.

***Supplementary Table 1*.** Gene set enrichment analysis (GSEA) on age related gene expression in morpahlino and control animals. 3733 orthologous human genes from 75212 probsets. Gene probe sets collapsed to gene summary by average. Group ANOVA by age groups. q value is the false discovery rate (fdr) modified p-value of the GSEA, ES = edge score, NES= normalised edge score, abs(ES)= absolute edge score. GSEA performed in Qlucore Omics Explorer (v3.6) using Gene Ontology Biological Process. Significance level of the fdr coloured from blue (highly significant) to red (not significant).

***Supplementary Table 2*.** Gene sets from the wider connected transcriptome of the 20-30dpf Control and Morpholino Zebrafish (460 and 12775 respectively). Age rank regression analysis of gene expression presented with R-statistic and p-value. Zebrafish gene symbol and corresponding human orthologue shown.

***Supplementary Table 3*.** A Gene probe sets (3460) from the wider connected transcriptome of 20-30dpf Morpholino Zebrafish dysregulated in controls. Age rank regression analysis of gene expression presented with p-value and q-value (false discovery rate modified p-value). ZF gene symbol and corresponding human orthologue shown.

***Supplementary Table 4.*** Dysregulated expression of cardiac phenotype marker genes in the KD zebrafish. False discovery rate modified p-value (q-value) shown for age group ANOVA.

---

**Supplementary Figure 1:**
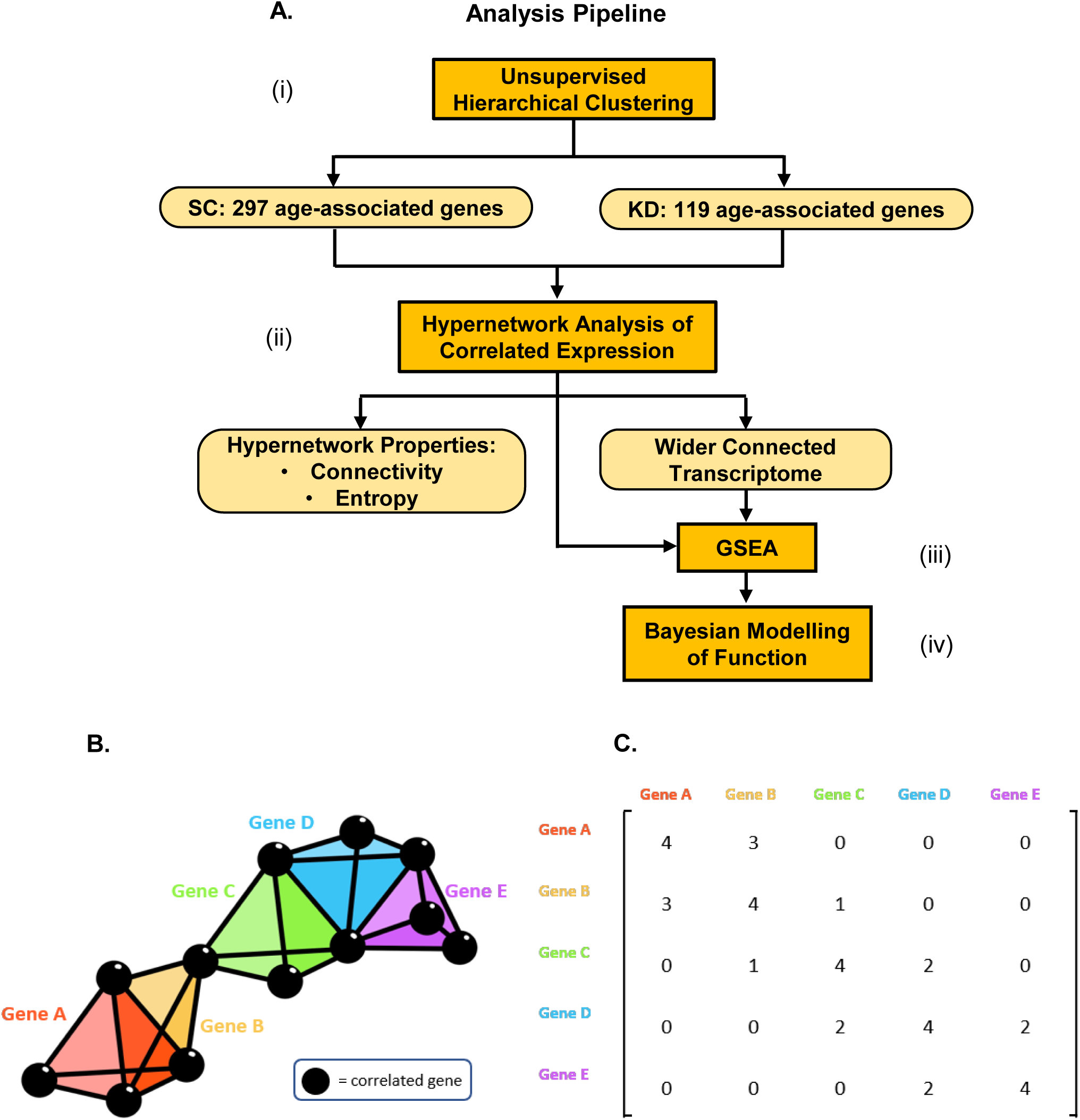
Pipeline of transcriptomic analysis and hypernetworks. ***1a.*** Analysis pipeline of transcriptomic data. **(i)** Unsupervised hierarchical clustering was performed to identify age-associated gene clusters. Genes were filtered by variance, using a projection score to maximise the informativeness of the genes selected. Clusters of age- associated genes were identified for SC and KD animals. **(ii)** Hypernetworks were generated using age associated genes for each group. Hypernetwork structure was quantified using connectivity and entropy. Clusters of highly connected genes were identified, and a wider set of transcripts were implicated as important by identifying the complete subgraph between cluster nodes and edges in the hypernetwork incidence matrix. **(iii)** GSEA was used to investigate biological functions associated with genes clustered by the hypernetwork or implicated by the complete subgraph in the hypernetwork incidence matrix. **(iv)** Biological processes identified by GSEA were assessed for functional activity. Hypernetworks were iterated, using subsets of genes associated with each process, and calculating hypernetwork entropy. A Bayesian modelling approach was used to detect differences in entropy distributions between processes. **1*b.*** A general model of a hypernetwork, shown as a three-dimensional representation of genes (coloured tetrahedra) correlating with the expression of other genes (black spheres). Shared correlations are represented by matching vertices, edges, and faces of the tetrahedra. The dimensionality of the connection between genes is defined by the number of shared correlations between those genes. ***1c.*** A hypernetwork representation of the “higher order” interactions within the transcriptome. This summary of the genes with shared correlations can be considered as the incidence matrix of a multi- dimensional network.

**Supplementary Figure 2.**
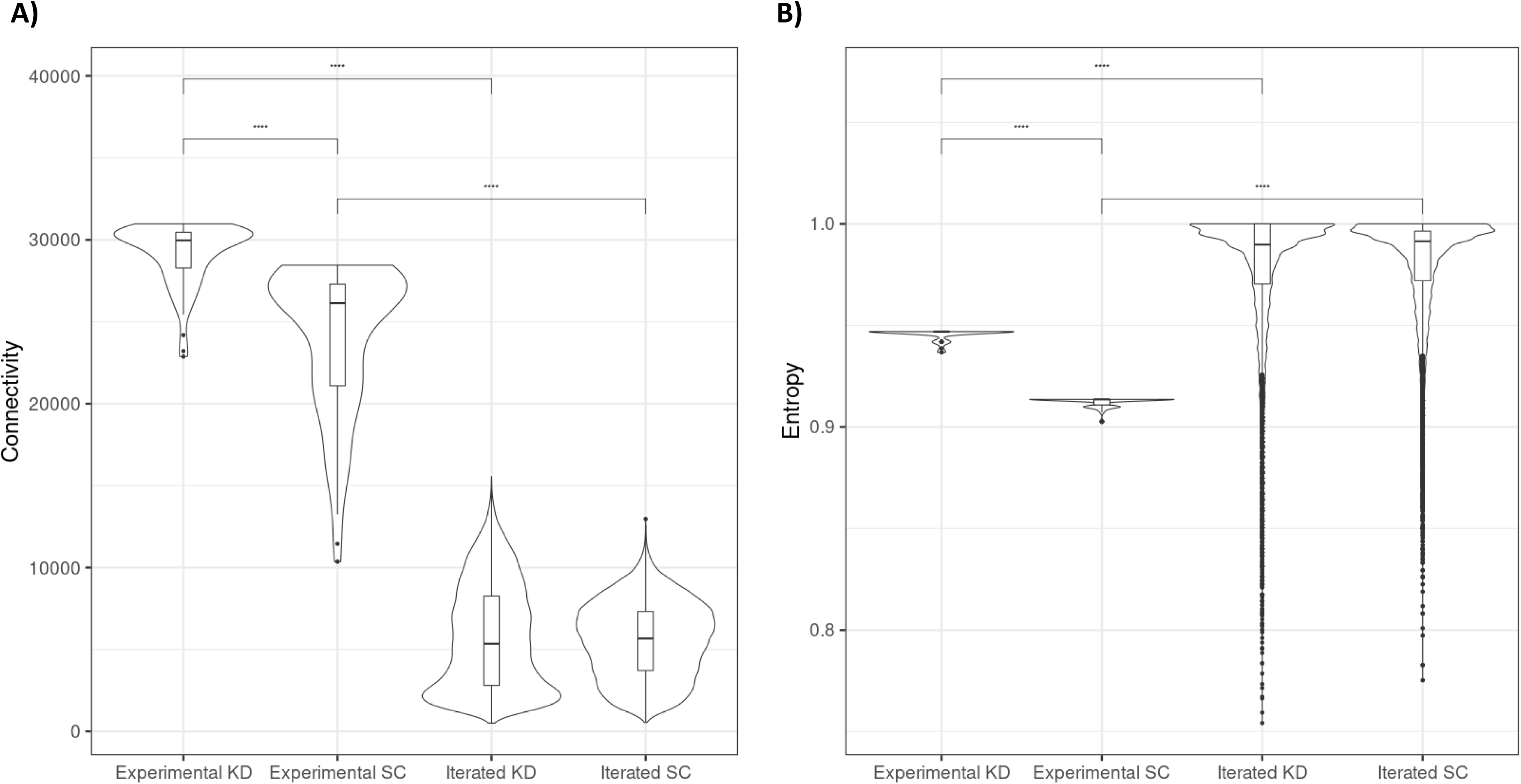
Connectivity (A) and entropy (B) of 20-30 dpf associated genes in the SC and KD zebrafish compared with randomly selected gene sets. 2*A.* Connectivity in the experimental data was significantly greater than in the random iterative data, and greater in the KD data than SC (p < 0.0001). 2*B.* Entropy was significantly lower in the experimental data than the random iterative data. The reduction in entropy between the experimental data and interated data was greater in the SC than the KD.

